# Neural Dynamics and Seizure Correlations: Insights from Neural Mass Models in a Tetanus Toxin Rat Model of Epilepsy

**DOI:** 10.1101/2024.01.15.575784

**Authors:** Parvin Zarei Eskikand, Artemio Soto-Breceda, Mark J. Cook, Anthony N. Burkitt, David B. Grayden

## Abstract

This study focuses on the use of a neural mass model to investigate potential relationships between functional connectivity and seizure frequency in epilepsy. We fitted a three-layer neural mass model of a cortical column to intracranial EEG (iEEG) data from a Tetanus Toxin rat model of epilepsy, which also included responses to periodic electrical stimulation. Our results show that some of the connectivity weights between different neural populations correlate significantly with the number of seizures each day, offering valuable insights into the dynamics of neural circuits during epileptogenesis. We also simulated single-pulse electrical stimulation of the neuronal populations to observe their responses after the connectivity weights were optimized to fit background (non-seizure) EEG data. The recovery time, defined as the time from stimulation until the membrane potential returns to baseline, was measured as a representation of the critical slowing down phenomenon observed in nonlinear systems operating near a bifurcation boundary. The results revealed that recovery times in the responses of the computational model fitted to the EEG data were longer during 5 min periods preceding seizures compared to 1 hr before seizures in four out of six rats. Analysis of the iEEG recorded in response to electrical stimulation revealed results similar to the computational model in four out of six rats. This study supports the potential use of this computational model as a model-based biomarker for seizure prediction when direct electrical stimulation to the brain is not feasible.

## Introduction

Epilepsy is a chronic neurological condition characterized by the occurrence of repetitive seizures arising from atypical neuronal activity in the brain [23]. Despite extensive research, the underlying mechanisms that contribute to the development and progression of epilepsy are still not understood. Unraveling the mechanism of epileptogenesis will assist to better understand the circuitry of the neurons in transition from normal to seizure state. This improved understanding of the complex mechanisms of epilepsy will aid in identifying novel therapeutic targets for treating this condition.

A compelling hypothesis posits that the shift in brain states from normal to seizure corresponds to a critical transition, marked by the presence of a critical threshold or tipping point, where such a transition occurs [7, 14, 15]. In dynamical systems theory, an important indicator of the system approaching a tipping point is the critical slowing down phenomenon [24]. This refers to the property of a nonlinear system in which it takes longer to recover from a perturbation as it approaches a critical point. A recent study has shown that EEG recordings exhibit signs of critical slowing down prior to seizures in epileptic patients, and suggests it as a biomarker for brain excitability [18].

One method to assess critical slowing down, and thus brain excitability, is “probing”. Probing involves applying a short pulse of electrical stimulation to the system and recording the response of the brain to the stimulation, which offers insights into the system’s behaviour through characteristics such as amplitude and temporal dynamics. Using probing to measure brain excitability is an effective approach for studying the pre-ictal period (the period of time immediately preceeding a seizure) [2].

While probing proves to be an excellent method for studying the pre-ictal period, it is not always feasible or practical to apply direct stimulation to the brains of patients. Recognizing these limitations, we turn to computational models as an indispensable tool in advancing our understanding of brain dynamics. In instances where direct probing may pose challenges, computational models offer an alternative avenue. By fitting background EEG data recorded from the brain to our computational models, we can simulate and study the response to probing in a computational model. This approach also allows us to gain valuable insights into the intricate dynamics of the pre-ictal period.

Here, we used intracranial EEG data recorded from a Tetanus Toxin (TT) rat model to fit the parameters of a neural mass model of a cortical column. TT has been shown to induce ongoing spontaneous seizures when injected into the hippocampus of rodents [6].The frequencies of resulting seizures wax and wane over a period of around 6 weeks. The exact mechanism of action of TT is still unknown. Some studies show that the TT inhibits the release of neurotransmitters from inhibitory neurons. However, excitatory transmission is also reduced ultimately until it is completely blocked [3, 12, 19]. Our aim was to investigate how functional connectivity correlates with seizure frequency.

We have investigated changes in the model parameters fitted to recorded background EEG data in response to varying numbers of seizures induced by the effects of TT. We also compared the results of the data driven model with the results of analysing the intracranial EEG data recorded in response to probing in rats. This comprehensive approach promises to shed light on the complex interplay between brain excitability, critical transitions, and the dynamics of epilepsy.

## Materials and methods

### Neural mass model

We have developed a neural mass model of a cortical column that comprises three interconnected motifs representing layer 2/3, layer 4 and layer 5 of the cortex [11]. Each motif in the neural mass model consists of one population of excitatory neurons and one population of inhibitory neurons, interconnected through forward and recurrent inhibitory and excitatory connections as shown in Fig. 1. Excitatory populations receive excitatory recurrent inputs and inhibitory inputs from within the same motif. Likewise, inhibitory populations receive inhibitory recurrent inputs and excitatory inputs from within the same motif (Fig. 1). Finally, both excitatory and inhibitory populations receive inputs from other motifs in the network. Additionally, external excitatory inputs from outside of the column, representing thalamic and inter-cortical connections, contribute to the overall neural dynamics.

**Fig 1.**
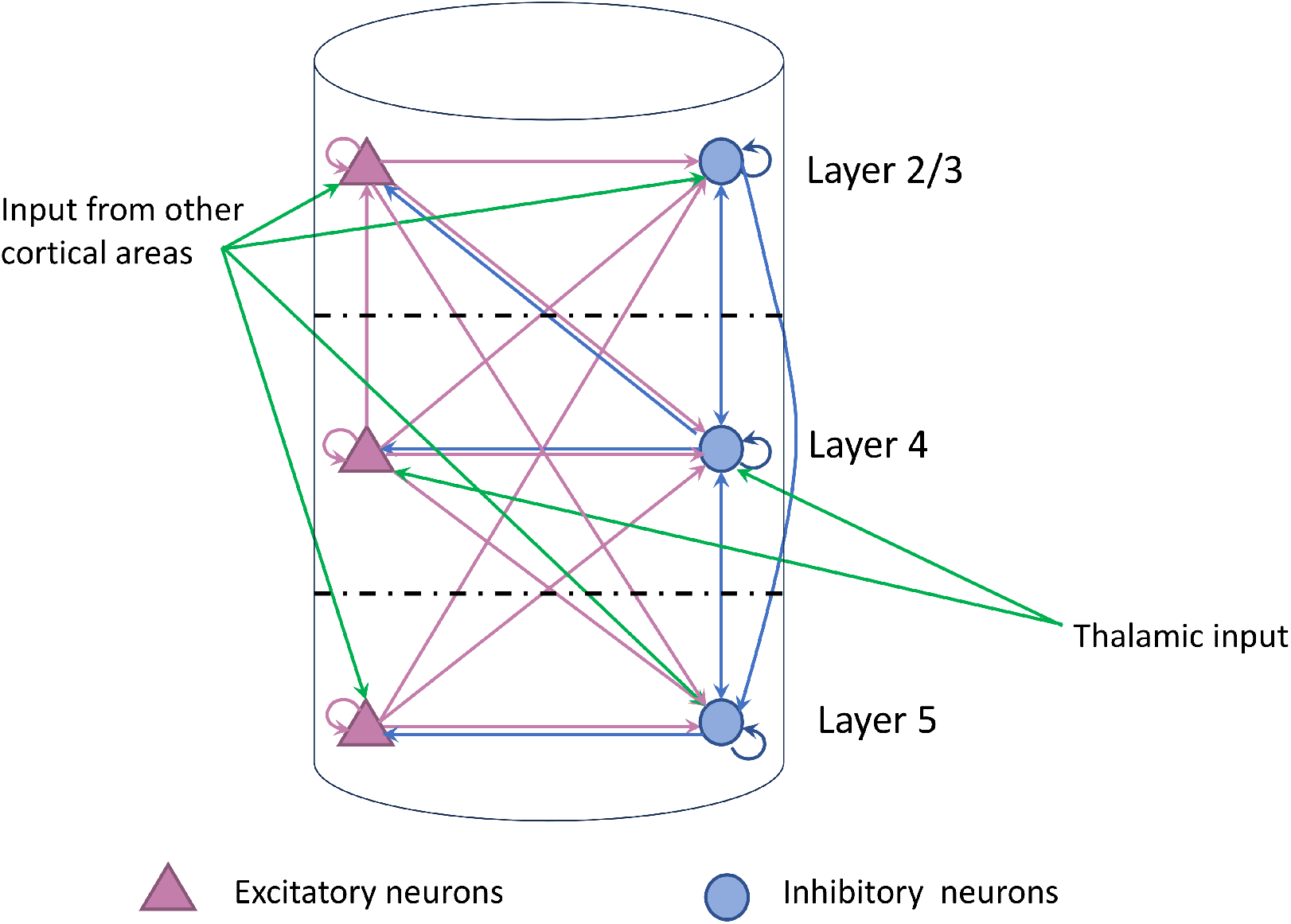
The structure of the model of the cortical column. The model comprises populations of excitatory neurons (purple triangles) and inhibitory neurons (blue circles), in layers 2/3, 4, and 5 of the cortical column. There are Excitatory and inhibitory connections between populations within the column, denoted by purple and blue arrows, respectively. Green arrows signify external input from other cortical areas and the thalamus

The neural populations are modeled as the current-based synapses with dynamics given by

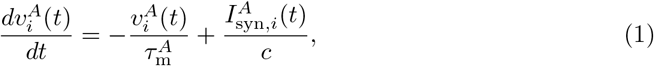

where 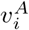 represents the mean membrane potential of a population of neurons, with superscript *A* indicating either excitatory (*E*) or inhibitory populations (*I*), and the subscript *i* indicating the layer number (2/3, 4, or 5). The parameter values are displayed in Table 1 [21]. The passive membrane time constants,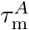, exhibit distinct values for excitatory and inhibitory neurons, and *c* indicates the membrane capacitance. The weighted summation of the incoming currents is given by

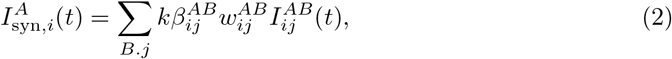

where 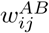 denotes the connectivity weights originating from population *B* in layer *j* to population *A* in layer *i*, as illustrated in Table 2, and have been derived from values obtained by the Allen Institute [1, 4, 11]. The values of 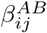 are computed by fitting the recorded EEG data to the output of the model. *k* acts as a correction factor, aligning values from single-cell recordings with the correct operational parameters for neural mass models, while preserving their relative strengths [11].

**Table 1.** Parameters of the model.

| Parameter | Description | Value |
| --- | --- | --- |
| $\tau_m^E$ | Membrane time constant of the excitatory neurons | 0.02s [21] |
| $\tau_m^I$ | Membrane time constant of the inhibitory neurons | 0.04s [21] |
| $c$ | Membrane capacitance | 0.35nF [10] |
| $\tau_s^{EE}$ | Synaptic time constant for excitatory-to-excitatory synapses | 8.5 ms [1, 4] |
| $\tau_s^{IE}$ | Synaptic time constant for excitatory-to-inhibitory synapses | 4.3 ms [1, 4] |
| $\tau_s^{EI}$ | Synaptic time constant for inhibitory-to-excitatory synapses | 13 ms [1, 4] |
| $\tau_s^{II}$ | Synaptic time constant for inhibitory-to-inhibitory synapses | 9 ms [1, 4] |
| $r$ | Gain (steepness) of the sigmoid function | 0.3 mV<sup>-1</sup> |
| $v_0$ | Postsynaptic potential for half of the maximum firing rate | 6 mV |
| $k$ | Connectivity constant | $5 \times 10^{-5}$ |
| $\phi_{\max}^E$ | maximum firing rate of excitatory neurons (where neurons saturate) | 21 Hz (inferred from [1]) |
| $\phi_{\max}^I$ | maximum firing rate of inhibitory neurons (where neurons saturate) | 30 Hz (inferred from [1]) |
| $w^E$ | Connectivity weight from other cortical areas/thalamus to excitatory neurons | 0.1 (inferred from [20]) |
| $w^I$ | Connectivity weight from other cortical areas/thalamus to inhibitory neurons | 0.2 (inferred from [20]) |
| $R_e$ | Membrane resistance | 100MΩ |

**Table 2.** Connectivity weights. The connectivity weights among neuronal populations in the model: ‘E’ denotes population of excitatory neurons, while ‘I’ designates population of inhibitory neurons spanning layers 2/3, 4, and 5. in layers 2/3, 4 and 5.

|  |  | Postsynaptic |  |  |  |  |  |
| --- | --- | --- | --- | --- | --- | --- | --- |
| Presynaptic |  | E2/3 | I2/3 | E4 | I4 | E5 | I5 |
|  | E2/3 | 0.06 | 0.88 | 0.01 | 0.25 | 0.06 | 0.16 |
|  | I2/3 | 0.35 | 1.06 | 0.06 | 0.18 | 0.02 | 0.06 |
|  | E4 | 0.11 | 0.21 | 0.2 | 1.14 | 0.07 | 0.24 |
|  | I4 | 0.29 | 0.59 | 0.48 | 1.46 | 0.07 | 0.17 |
|  | E5 | 0.01 | 0.13 | 0 | 0.13 | 0.09 | 0.27 |
|  | I5 | 0.04 | 0.06 | 0.02 | 0.11 | 0.49 | 1.51 |

The default connectivity weights in our modeled are calculated by considering connection probabilities and synaptic strengths given in Synaptic Physiology database (Allen Institute, USA) [4, 25] by multiplying their associated values.Details of the methods of deriving the weights are given in [11]. The connectivity weights pertaining to intra-columnar connections are detailed in Table 2.

The dynamics of post-synaptic currents in the model are modelled using an exponentially decaying function [5, 8],

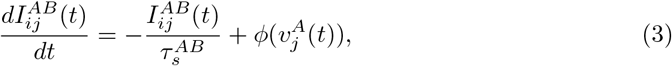

The synaptic time constants 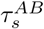 are adapted from a recent computational investigation conducted by Billeh et al. [4]. The output firing rate of a neural population is determined by the function *ϕ*(), which is derived by applying a sigmoid function to the membrane potentia,

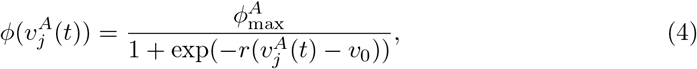

Here, 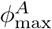 represents the peak firing rate of the neural population, *v*_0_ denotes the postsynaptic potential at which the firing rate achieves half of its maximum value, and *r* governs the steepness or gain of the firing rate function.

### State and parameter estimation

The output EEG signal is modelled as the summation of the dipole currents (i.e., excitatory to excitatory and inhibitory to excitatory synaptic currents of all cortical layers) multiplied by the value of the membrane resistance. The observation equation has the following form

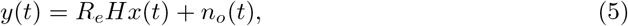

where *y*(*t*) is the modeled EEG and *R*_*e*_ is a scaling constant. *H* is the observation vector containing values of 0, 1 and −1. A value of 1 is associated to an excitatory input current, a value of −1 is associated to inhibitory input current and a value of 0 refers to a synapses where the post-synaptic population is not excitatory. *n*_*o*_(*t*) ∼*N* (0, *R*) is the observation noise with the variance of *R*.

The aim is to estimate the parameters 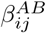 of the model and the states (i.e., the membrane potentials and synaptic currents) by fitting the output of the model to the EEG signals recorded from rats in control and TT groups. 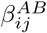 parameters scale the default values of connectivity weights given in Table 2. ℬ is a vector containing all of the 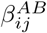 parameters that needs to be estimated.

One of the challenges in parameter estimation using an optimization algorithm is the absence of a unique solution. Consequently, the algorithm yields thousands of solutions. To address this issue, we define parameter dynamics in a manner that constrains the optimization solutions to an area closer to the biological ranges of parameters obtained in neurophysiological experiments. The dynamics for the parameters are modelled as

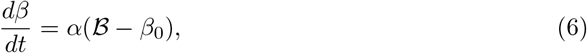

where *β*_0_ is a vector of ones that acts as a fixed-point to restore the values of 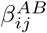 towards 1 with a decay constant of *α*. These dynamics were introduced to constrain the estimated parameters to be close to the default connectivity weight while minimizing the error of the estimation at the output. This technique also prevents parameters from drifting over time.

For simultaneous estimation of parameters 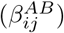 and states, we put them together in a single augmented vector, *x*. As a result, the equation defining the model in matrix notation is

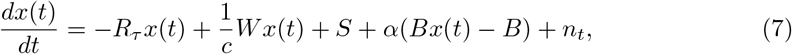

where *x* is the augmented state vector representing membrane potentials, synaptic currents and 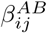 parameters that are to be estimated. *R*_*τ*_ is a vector with membrane time constants and synaptic time constants in the elements corresponding to states in *x*, and values of 0 in the elements corresponding to 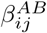 parameters in *x*, and takes the form

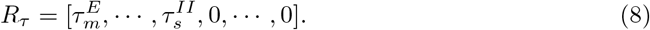

*W* is a matrix containing the default values for the connectivity weights multiplied by 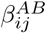 parameters as the scale of these default values; the other elements of the matrix that are associated with dynamics of the synaptic currents and also the dynamic of the *β* parameters are 0:

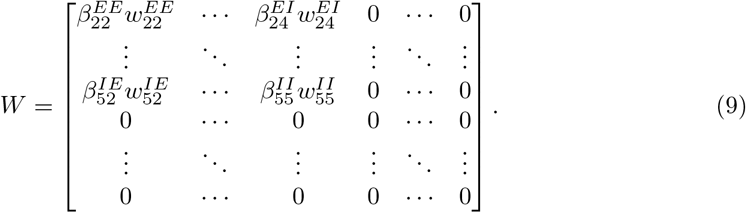

*S* is a vector with the elements of the sigmoid functions associated with the dynamics of the current in the state vector, and elements of 0 that are associated with the dynamics of parameters of 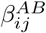 and the membrane potentials,

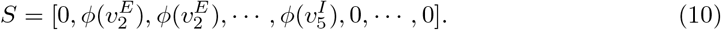

*B* is a vector of 0’s and 1’s in the form of

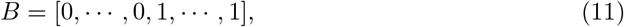

where values of 1 are associated with the 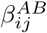 parameters and 0 values are associated with other elements in the state vector, *x. n*_*t*_ is a vector for model noise with the variance of *Q n*_*t*_ ∼ *N* (0, *Q*).

To estimate the parameters and states of the model, we use the Unscented Kalman Filter (UKF)[13]. The filter estimates the values of parameters and states to achieve the minimum mean squared error estimates of the output of the nonlinear model compared to the recorded EEG signals. The goal of this method is to compute the most likely posterior distribution (*a posteriori* state estimate) conditioned on the previous observation signal (i.e., recorded EEG signals) with the *a posteriori* mean and covariance of

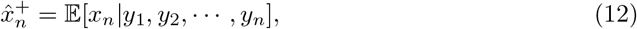

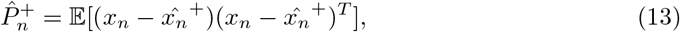

where *n* is the index in discrete time notation.

The estimation process has two stages. The first stage predicts where the prior distribution, obtained from the previous estimate, is propagated through the equations of the model. The outcome of this step is called the *a priori* estimate. The mean and covariance of the *a priori* estimate are

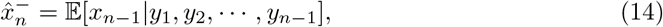

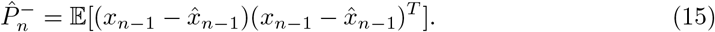

The weights to correct the *a priori* estimate is called Kalman gain,*K*, and is defined as

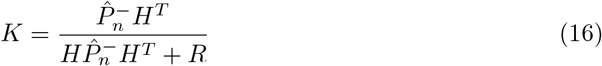

As the system is nonlinear, the integration required for the estimation of the expectations is carried out through an approximate solution. The Unscented Transform approximates the statistics of the random variables (with mean and covariance 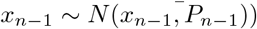, that undergo the nonlinear transformation. As the first step, we define the sigma vectors, which are put together in the matrix,χ,

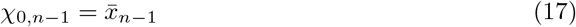

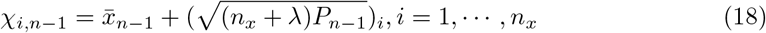

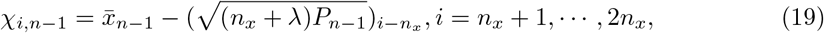

where (2*n*_*x*_ + 1) is the number of the sigma vectors and *λ* = *γ*^2^(*n*_*x*_ + *κ*) *™ n*_*x*_ is a scaling parameter with *γ* a constant with a positive value of 1 that determines the spread of the sigma points around their mean, 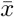, and *κ* is a secondary scaling parameter that is usually set to 3 or 4. The resulting sigma vectors are propagated through the nonlinear model,

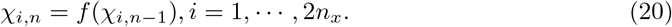

The mean and covariance of the transformed sigma vectors (i.e., Eq.14 and Eq.15) after undergoing the nonlinear transformation of the model are approximated as

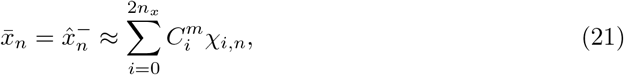

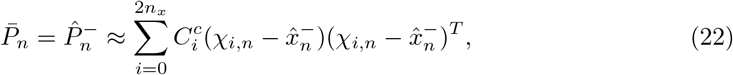

where the weights of the sigma vectors, *C*_*i*_, are computed using

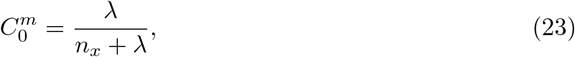

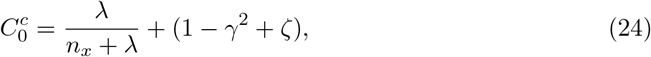

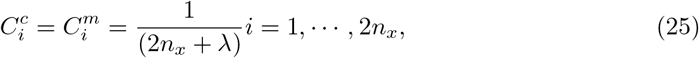

where ζ is a constant with the value of 2.

The second stage of the estimation process updates the parameters to shift the estimated posterior depending on the recorded EEG signals. As the observation equation, Eq.5, is linear, we can use the standard Kalman filter Bayesian update equations

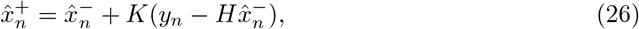

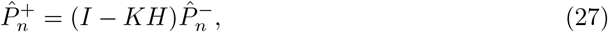

where *I* is an identity matrix. After each step, the *a posteriori* estimate is replaced with *a priori* estimate [13].The time step for the Unscented Kalman Filter is 1/2048 s.

After the initial round of optimization, the resulting estimated parameters are replaced with the default connectivity weight for the second round of estimation. This process enables the alteration of the restricted solution region, aligning it with a new set of values that more accurately reflect the connectivity strength estimated for individual rats.

### Data

The data for this study was collected from a intra-hippocampal TT rat model study [6]. Briefly, six rats were placed in the TT group after intra-hippocampal injection of TT and four rats were placed in the control group after instead administrating phosphate-buffered saline. Five stainless steel electrodes were implanted into the skull of each rat for intracranial EEG recording. After around 1-2 weeks, the TT rats started to have spontaneous seizures. The daily numbers of seizures waxed and waned over a period of approximately 6 weeks, as illustrated in Fig. 2 for one TT rat. Between 4 to 5 weeks after injection, the seizure rate tended to decrease until there were no seizures detected. EEG data were recorded continuously at 2048 Hz sampling rate for 23 hours every day, with 1 hour for daily maintenance checks and back up of the data. Six rats are used from the original data as one of the rats died on Day 26.

**Fig 2.**
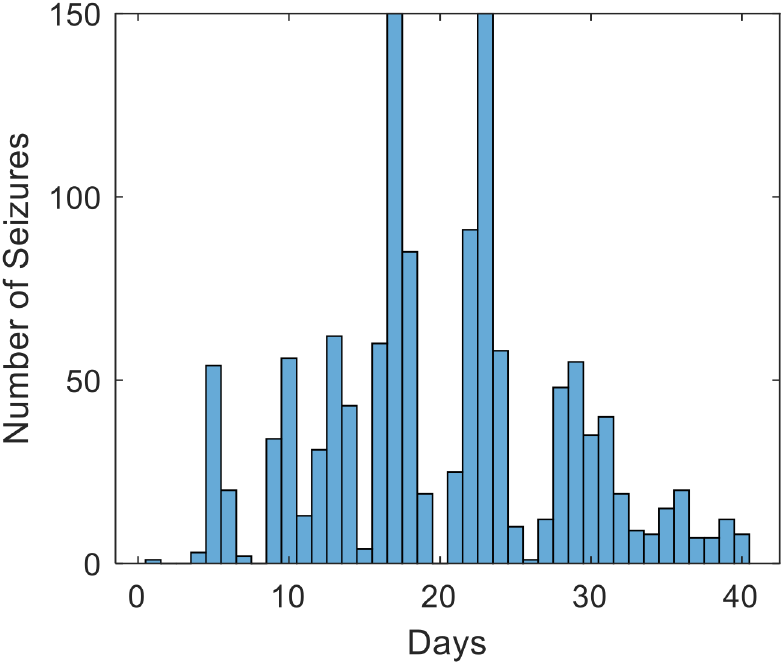
Seizure occurrences over time for an example TT rat. The x-axis is days since the start of the recording. The y-axis is the total number of seizures that occurred on the specific day, regardless of whether they occur during the probing-on or probing-off periods.

The experimental protocol included periodic, low-level electrical stimulation or “probing” [6]. The probing-on period consisted of 100 biphasic pulses (pulse-width 0.5 ms and current/phase 1.26 *±* 0.024 mA) with inter-stimulus intervals of 3.01 s over a period of about 5 minutes. After each probing-on period, there was a 5 min probing-off period before the start of the next probing-on period. The research procedures received approval from the St. Vincent’s Hospital, Melbourne Animal Ethics Committee and were carried out following the guidelines outlined in the “Australian Code for the Care and Use of Animals for Scientific Purposes, 8th Edition” (2013).

### Fitting the Neural Mass Model to EEG Recordings and Comparative Analysis

The neural mass model was fitted exclusively to probing-off segments of the EEG recording during the non-seizure periods. Due to the random occurrence of seizures, the duration of EEG segments during probing-off periods may vary. On average, each segment lasts between 2 to 3 minutes. Initially, we fitted the model to the first 20 probing-off segments recorded each day. Fitting the entire daily recorded dataset to the model was computationally demanding, necessitating our decision to restrict the fitting to only the first 20 recordings of each day. We calculated the averages of the parameters estimated from 20 recordings for each day, for each of the six rats in the TT group. We then calculated the correlations between these parameters and the trend in the daily changes in the number of seizures for each rat. To obtain this trend in seizure changes, we applied a five-day moving average to the seizure counts. Subsequently, we performed linear regression analysis and generated model statistics, reporting the p-value and F-value of the correlation.

Then, we fitted the neural mass model to background EEG recordings during preictal (5 minutes before seizures) and 1 hour before seizures. The background EEG signal refers to the probing-off sections of EEG recordings where no seizures have occurred. We considered only the lead seizures where there had not been any seizures at least 1.5 hours before these lead seizures. A 1 min duration of each background EEG recording from the sections within 5 mins before lead seizures and 1 hour before lead seizures are selected to fit the neural mass model. This comparison allowed us to assess the model’s behavior distant from and close to the times of seizures. The results of the modeling were compared with the response to stimulation in the data recorded during the stimulation probing-on periods.

We analyzed the EEG recordings in the probing-on sections, where 100 single-pulse electrical stimulation was applied every 3.01 s for periods of approximately 5 min. The waveform of the recorded response to one example of these probes is shown in Figure 8a. To investigate the critical slowing phenomenon, we measured the recovery time to stimulation as the time from the first peak in the waveform until the response returned to and remained within 10% of the baseline.To automatically detect where the waveform stabilizes at 10% of the baseline, the algorithm detects the last peak in the waveform and extracts the time when the waveform reaches 10% of the baseline, starting from the last peak. We measured the recovery time to stimulation of the 20 last stimuli of each probing-on period, as illustrated in Figure 8b. We calculated the average of the recovery times of these 20 probes.

## Results

We fitted the neural mass model to background EEG recordings segments from each day of recordings. Fig. 3a shows one EEG recording from rat T6 in the TT group. This figure shows how well the model fitted the original EEG recordings. The estimated EEG signal (blue) closely matches the original EEG signal (green), although there is a degree of mismatch between the results of the model and the original EEG during sections of EEG signal with more rapidly fluctuating activity. Fig. 3b, c and d show the membrane potentials and synaptic currents driven by the data in layers 2/3, 4 and 5.

**Fig 3.**
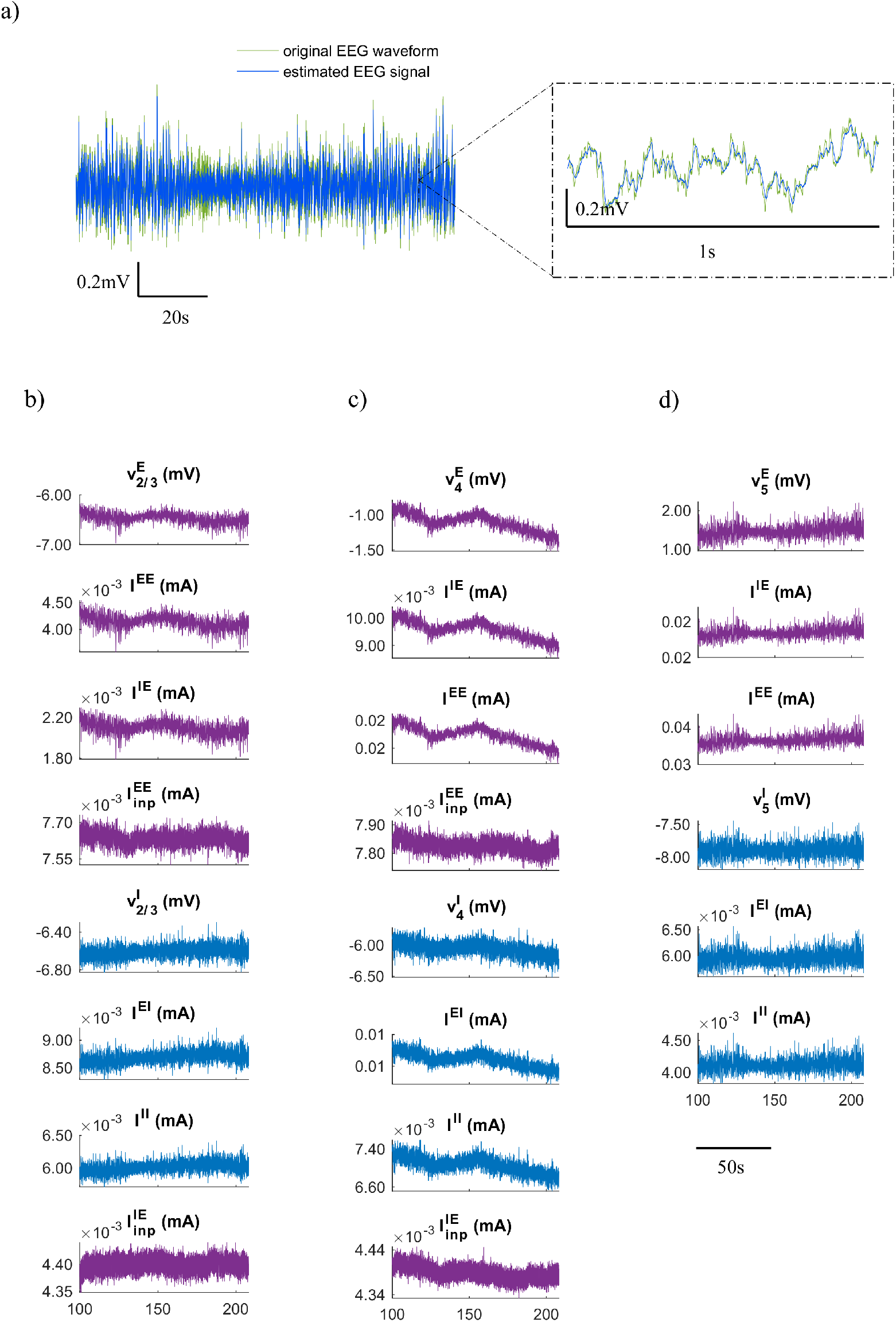
a) Estimated EEG signal (blue) compared to the original EEG waveform (green) recorded from the rat T6. b) Data driven membrane potentials of excitatory neurons and excitatory synaptic currents (purple) in layer 2/3 and membrane potentials of inhibitory neurons and inhibitory synaptic currents (blue) in layer 2/3. The heading on top of each graph indicates the membrane potential of excitatory 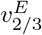 or inhibitory 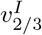 neurons in layer 2/3 or the synaptic currents across excitatory to excitatory neurons *I*^*EE*^, excitatory to inhibitory *I*^*IE*^, Inhibitory to excitatory *I*^*EI*^ or inhibitory to inhibitory *I*^*II*^ neurons. c) Data driven membrane potentials and synaptic currents of neural populations in layer 4. d) Data driven membrane potentials and synaptic currents of neural populations in layer 5.

Fig. 4 shows the estimated parameters of the model (*β*) after fitting the sample EEG recording in Fig. 3a. *β* the deviation of the connectivity weights from their default values. In this specific example, the connectivity weight of excitatory-to-excitatory connections in layer 5, 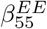, deviates the most from its default value, while the inhibitory-to-inhibitory connectivity weight from layer 5 to 4, 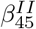, does not vary much at all.

**Fig 4.**
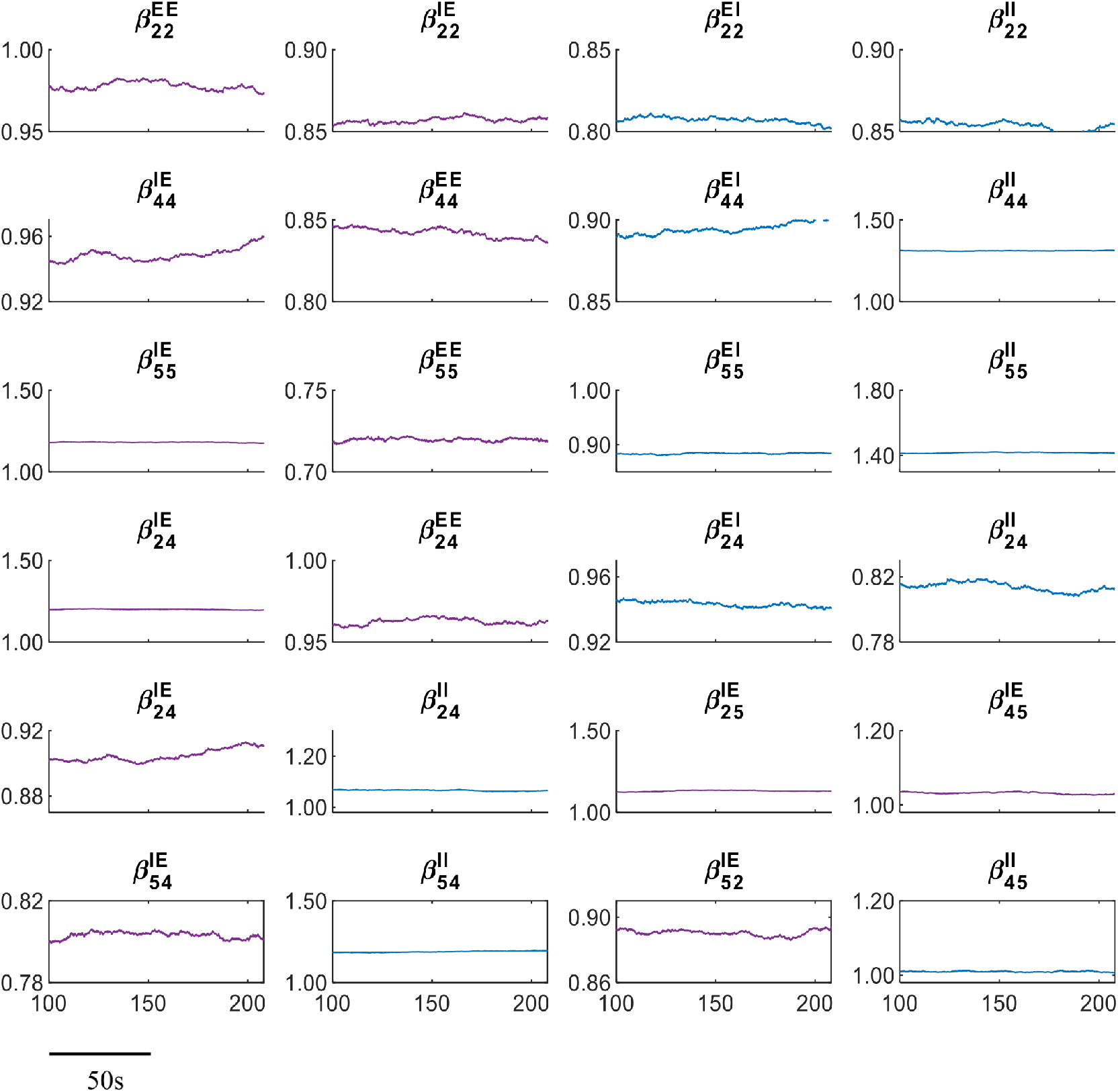
Data driven model parameters of *β* for excitatory connectivity weights (purple) and *β* for inhibitory connectivity weight (blue) fitted to one recording of the rat T6 (Fig. 3a). The closer the values of *β* to one, the more similar the connectivity weights are to the default values of connectivity weights.

### Correlation of the model parameters with the number of daily seizures

Fig. 5 and Fig. 6 show the parameters of the model that have statistically significant correlation with the number of daily seizures (i.e.. p-value less than 0.05 and F-value above the critical value of 4.12) in different rats. The parameters of the excitatory synapses are shown in Fig. 5 and the parameters of the inhibitory synapses are shown in Fig. 6.

**Fig 5.**
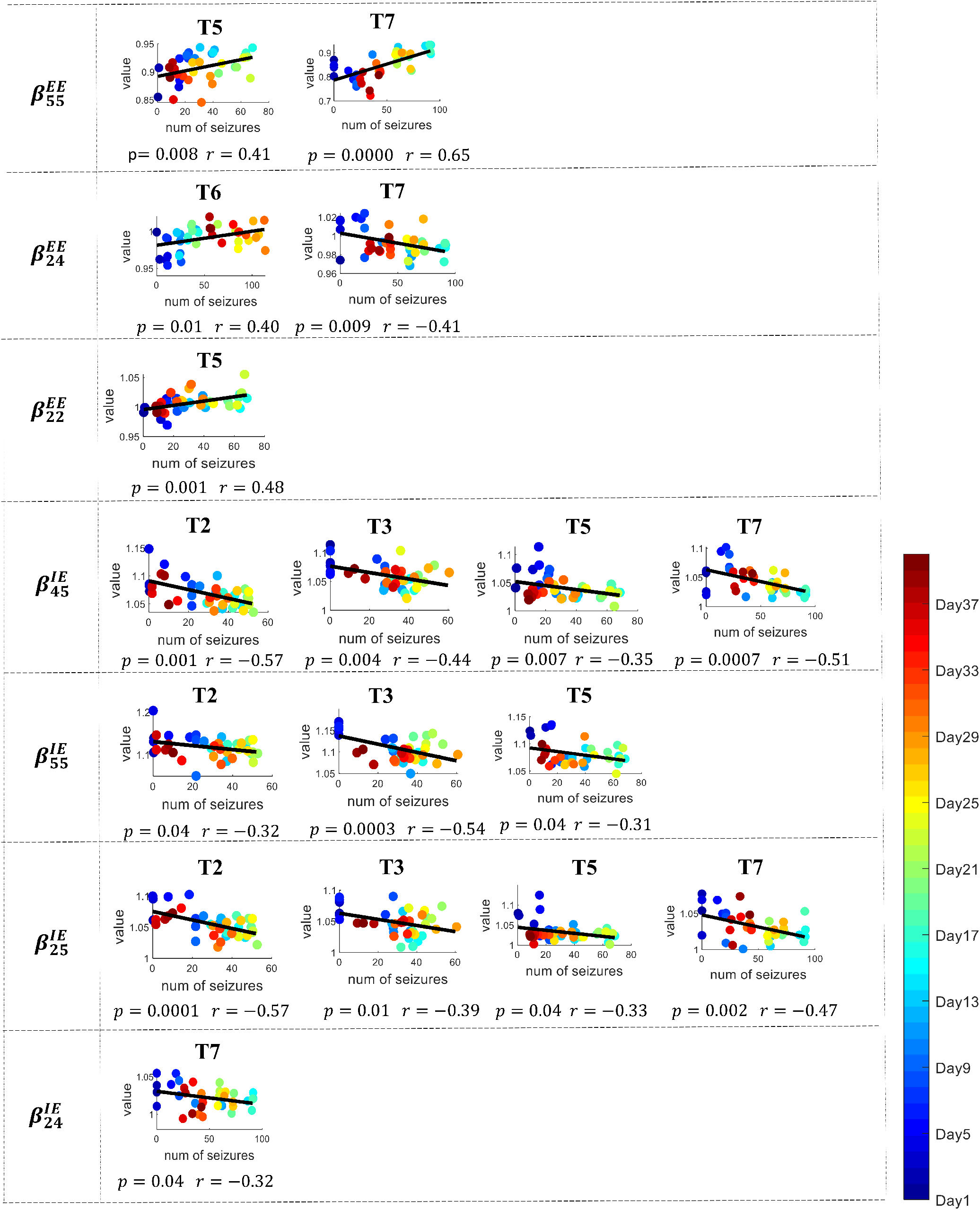
Average daily parameter estimates of excitatory synapses of the model that had significant correlations with daily seizure rates in different rats. Each dot represents the daily average of the parameter value and is plotted against the number of seizures that day. The parameter measured is shown on the left. On top of each panel is a label indicated which TT rat’s data is shown. The black solid line represents the fitted linear regression model. The p-value and r-value of the relationship between parameters and the number of the daily seizures are given below each graph. The colors indicate the progress over time from day 1 to day 40 of recordings.

**Fig 6.**
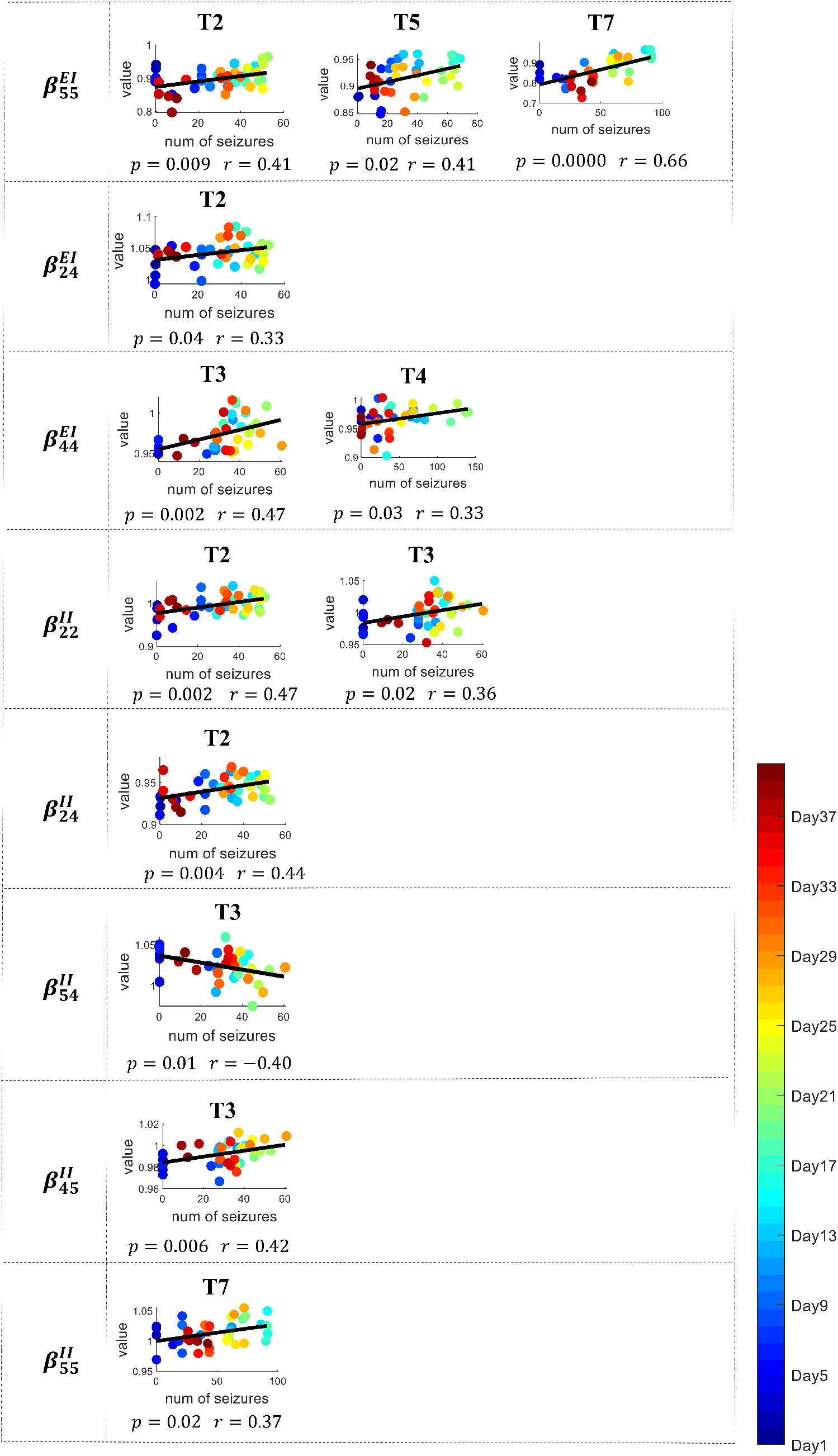
Average daily parameter estimate of inhibitory synapses of the model with significant correlation with daily seizures rate in different rats. Each dot represents the daily average of the parameter value compared to the number of daily seizures.The parameter symbol is shown on the left. The label of rats is shown on the top of each panel. The black solid line represents the best fitted linear regression model. The p-value and r-value of the relationship between parameters and the number of the daily seizures are given below each graph. The color bar shows the progress over time from day 1 (blue) to day 40 (red) of recordings.

Table 3 summarizes Figures 5 and 6, displaying the count of model parameters exhibiting significant positive or negative correlations (p-value less than 0.05) with the daily number of seizures across all rats. These results show that four out of five excitatory-to-excitatory connections had positive correlations with the numbers of daily seizures, indicating that the strength of excitatory-to-excitatory connections increased with increasing numbers of daily seizures. Six out of six inhibitory-to-excitatory connections (with statistically significant correlation) had positive correlations with the numbers of daily seizures and this is also the case for five out of six inhibitory-to-inhibitory connections. 12 out of 12 excitatory-to-inhibitory connections had negative correlation with the numbers of daily seizures, indicating that the strength of excitatory-to-inhibitory connections decreased with increasing numbers of daily seizures.

**Table 3.** Summary of the negative or positive significant correlation (p-value less than 0.05) of the parameters with the number of daily seizures across all rats.

| Connections | Number of connections | Increase strength | Decrease strength |
| --- | --- | --- | --- |
| Excitatory-to-Excitatory | 5 | 4 (80%) | 1 (20%) |
| Inhibitory-to-Excitatory | 6 | 6 (100%) | 0 (0%) |
| Inhibitory-to-Inhibitory | 6 | 5 (83%) | 1 (17%) |
| Excitatory-to-Inhibitory | 12 | 0 (0%) | 12 (100%) |

Figure 7 presents the correlation coefficients between trends in seizure counts and various parameters in the model. It also displays the critical correlation values (positive and negative) corresponding to significance level of 0.05 after adjustment for multiple comparisons using the Benjamini-Hochberg method. The r values corresponding to p-values larger than 0.5 are plotted in a faint colour. There is a general consistency in whether the parameters have positive or negative correlations with the number of seizures, regardless of the criteria used to establish the significant p-values.

**Fig 7.**
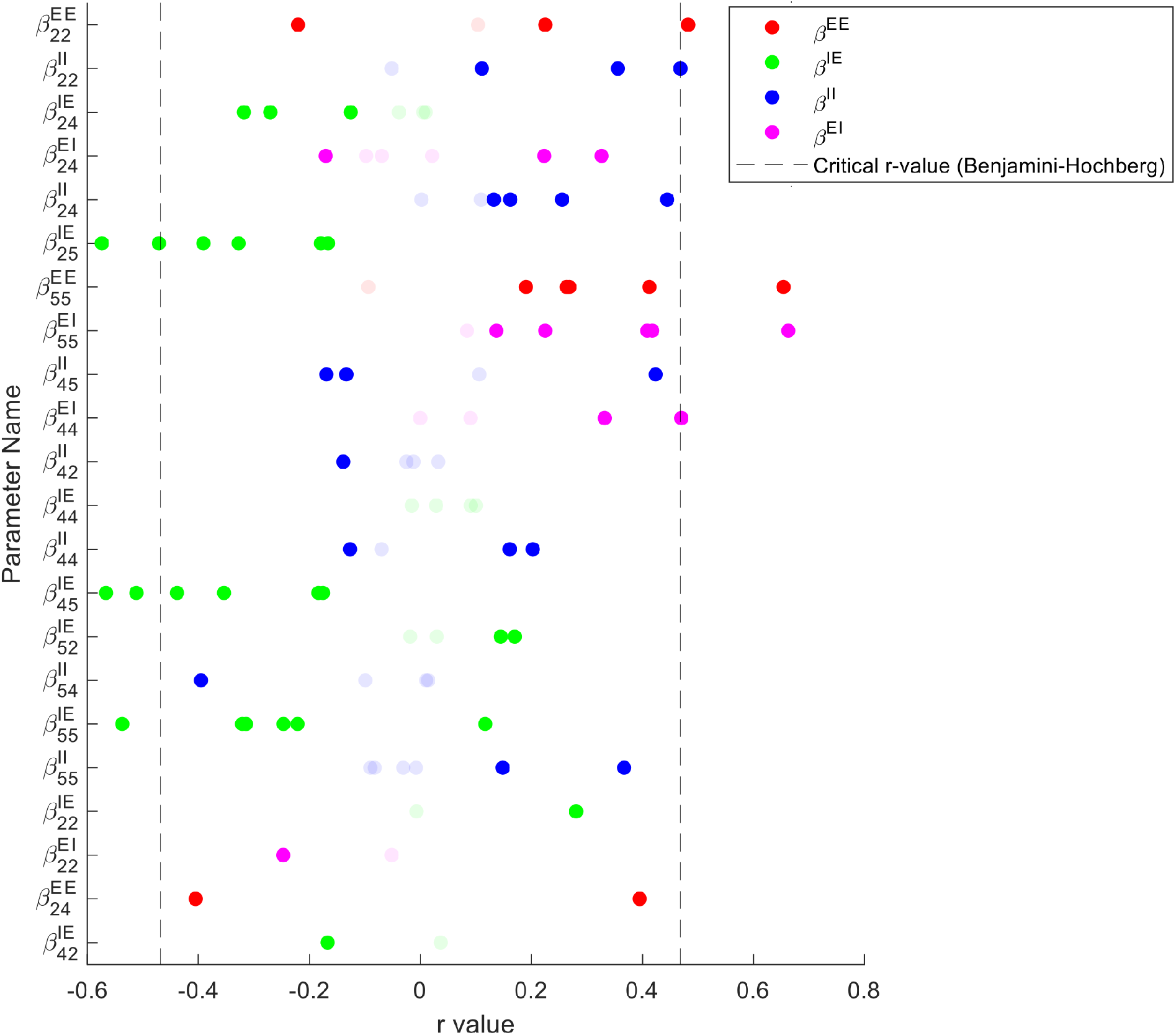
Scatter plot showing the correlation coefficients between trends in seizure counts and various parameters. Each dot represents the correlation coefficient for a specific parameter. The parameters are color-coded based on their category: red for parameters of excitatory to excitatory connections (*β*^*EE*^), green for excitatory to inhibitory connections (*β*^*IE*^), blue for inhibitory to inhibitory connections (*β*^*II*^), and purple for inhibitory to excitatory connections (*β*^*EI*^). Vertical lines indicate the positive and negative critical r-values corresponding to significance level of 0.05 after adjustment for multiple comparisons using the Benjamini-Hochberg method. The r values corresponding to p-values larger than 0.5 are plotted in a faint colour.

### Critical slowing down measure as an active biomarker Results from the data analysis

The average recovery time for probing periods within 5 mins before seizures compared to 1 hour before seizures across different rats is shown in Figure 8c. We considered only the lead seizures where there had not been any seizures at least 1.5 hours before these lead seizures. Red represents the average of the recovery time over all the sections within 1 hour before seizures. The error bar represents the confidence interval of this average over all seizures. Blue represents the average of the recovery time within 5 min before seizures. Across all of the rats, we removed 39 points out of 3060 points that had recovery time longer than 100 ms as the outliers from the analysis. Excessively longer durations are the result of changes in the baseline over time, leading to the failure of the algorithm.

The results show that the average recovery time 5 min before seizures was longer than the recovery time 1 hour before seizures for four rats. This was not the case for rat T4 and there is no significant difference observed for rat T5.

### Modeling results

Figure 9 shows the mean squared error, which entails averaging the squares of the differences between the estimated signals and the original EEG signals. After fitting the EEG recordings, we applied excitatory stimulation to both excitatory and inhibitory populations in different layers by delivering a pulse stimulus as the excitatory input from the thalamus and other cortical areas. The stimulation is applied to the population of neurons in all layers (layer 2/3, layer 4 and layer 5). The amplitude of the pulse stimulus is set at 0.5 mA with a duration of 100ms.

**Fig 8.**
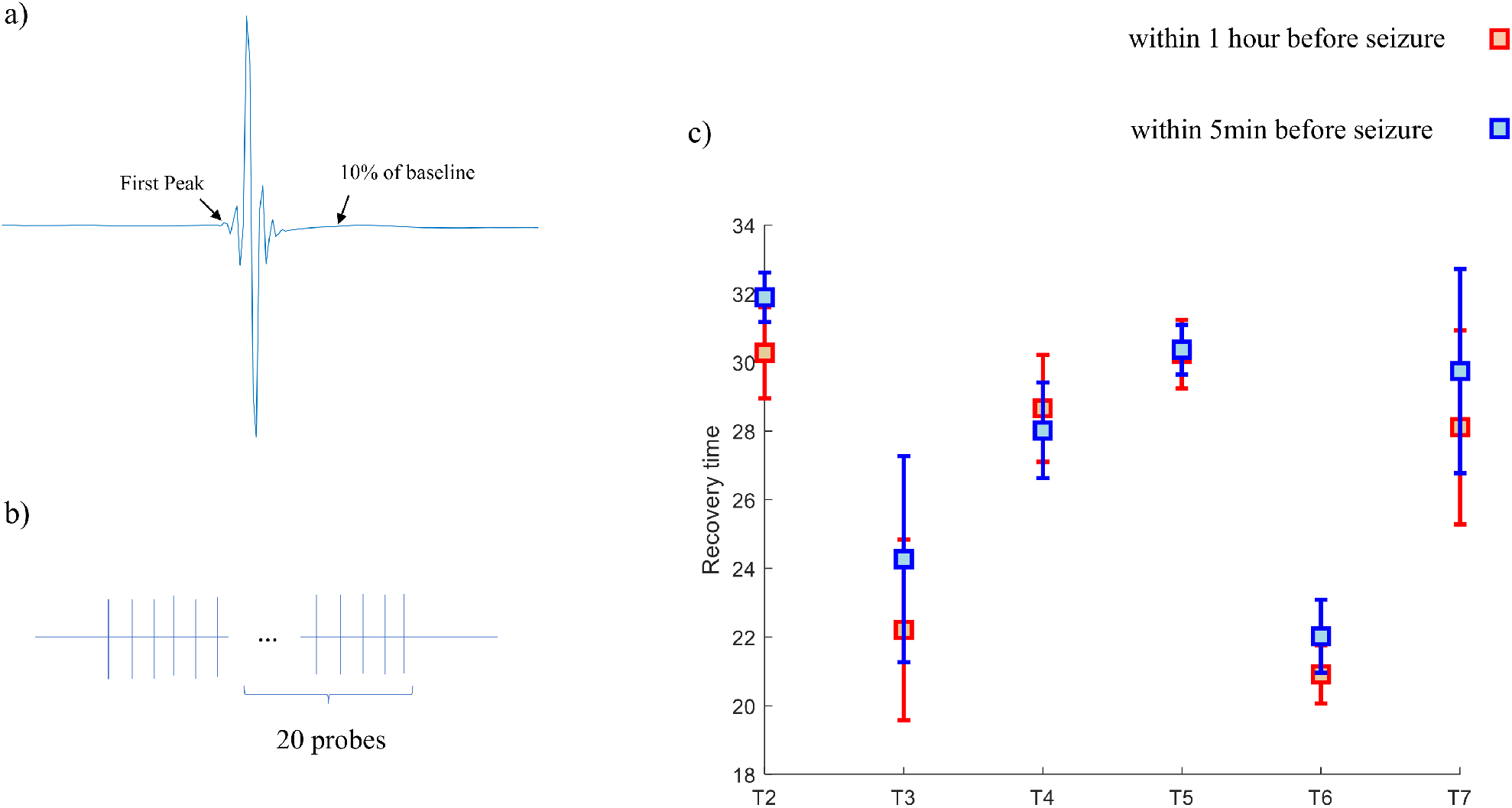
The recovery times of responses to probing in the EEG recordings. a) A sample waveform of the response to one probe in the data. We measured the recovery time from the stimulation by measuring the time of the first peak in the waveform until the response remained within 10% of the baseline. b) A schematic diagram of a probing-on section. We measured the average of the recovery time of the response to 20 last probes of each probing-on section. c) The average recovery time of the sections within 5 mins before seizures compared to 1 hour before seizures across different rats (T2, …, T7). The error bars represent the confidence intervals. Blue represents 5 mins before seizures and red represents 1 hour before seizures.

**Fig 9.**
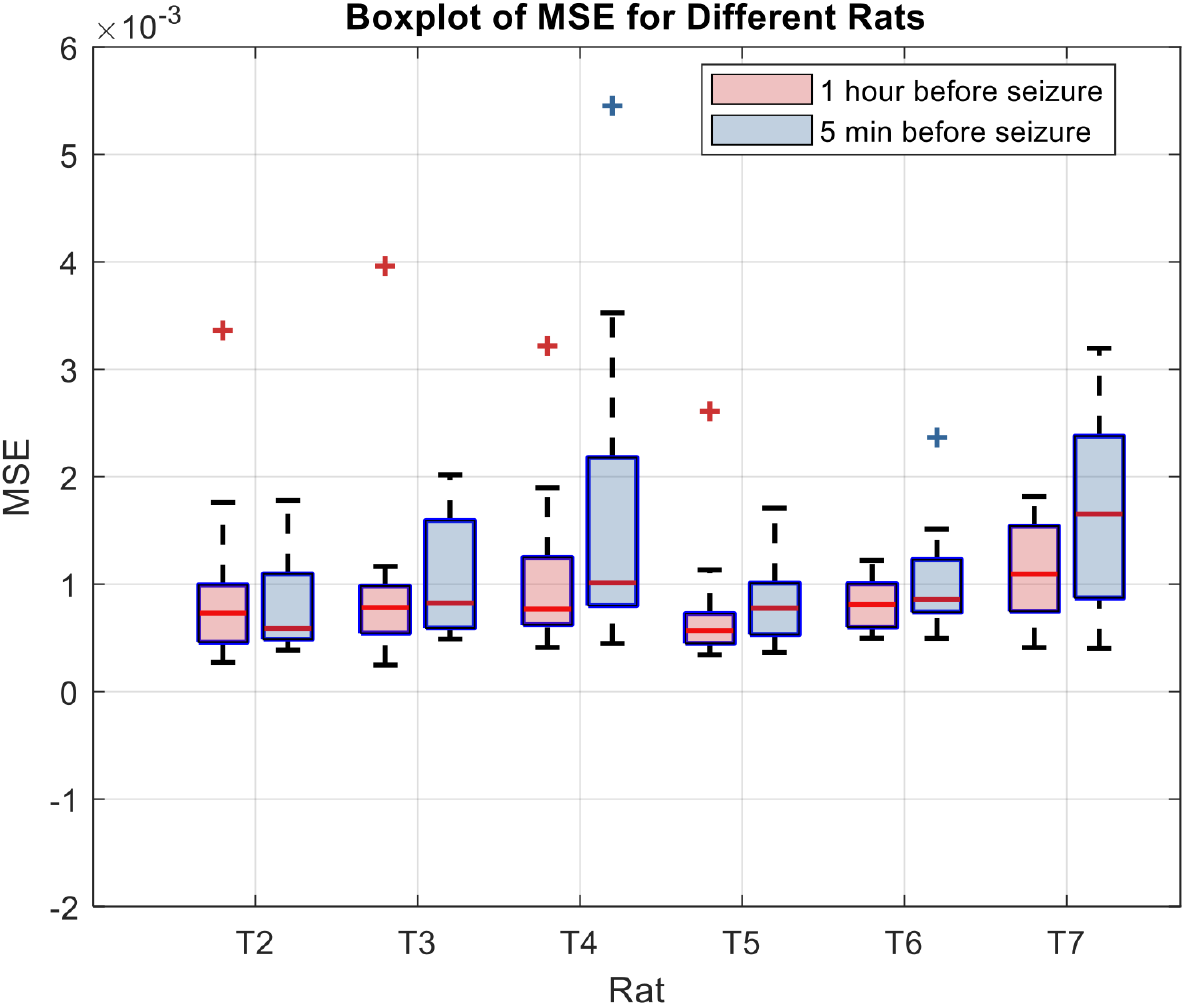
Boxplot visualization of mean squared error (MSE) in signal estimation for different rats. Each box represents the distribution of MSE values for two conditions: 1 hour before seizure (red) and 5 min before the seizure (red). Whiskers extend to 1.5 times the interquartile range, and outliers are displayed as individual points. The mid lines indicate the 95% confidence interval for the median. MSE values were computed across multiple signals for each rat.

Figure 10a shows an example of the response of excitatory neurons to stimulation. We measured the recovery time as the time starting from the beginning of stimulation until the response remains within 1% of the baseline. As there is no added noise in the model, we lowered the threshold from 10% to 1% of the baseline for more precise measurements. Figure 10b shows the measured recovery time for the model fitted to the background EEG signals 5 mins before seizures compared to 1 hour before seizures for TT rats T2-7. The error bars represent the confidence intervals. Blue represents 5 mins before seizures and red represents 1 hour before seizures. We excluded 2 out of 81 data points as outliers because they had unusually short recovery times below 250 ms when displaying the results. The results show that the average recovery time 5 mins before seizures was longer than 1 hour before seizures in four rats (T2, T3, T5, T7). However, this was not the case for two rats (T4, T6). The modeling results accord with the results from the EEG analysis in four rats (T2, T3, T4, T7).

**Fig 10.**
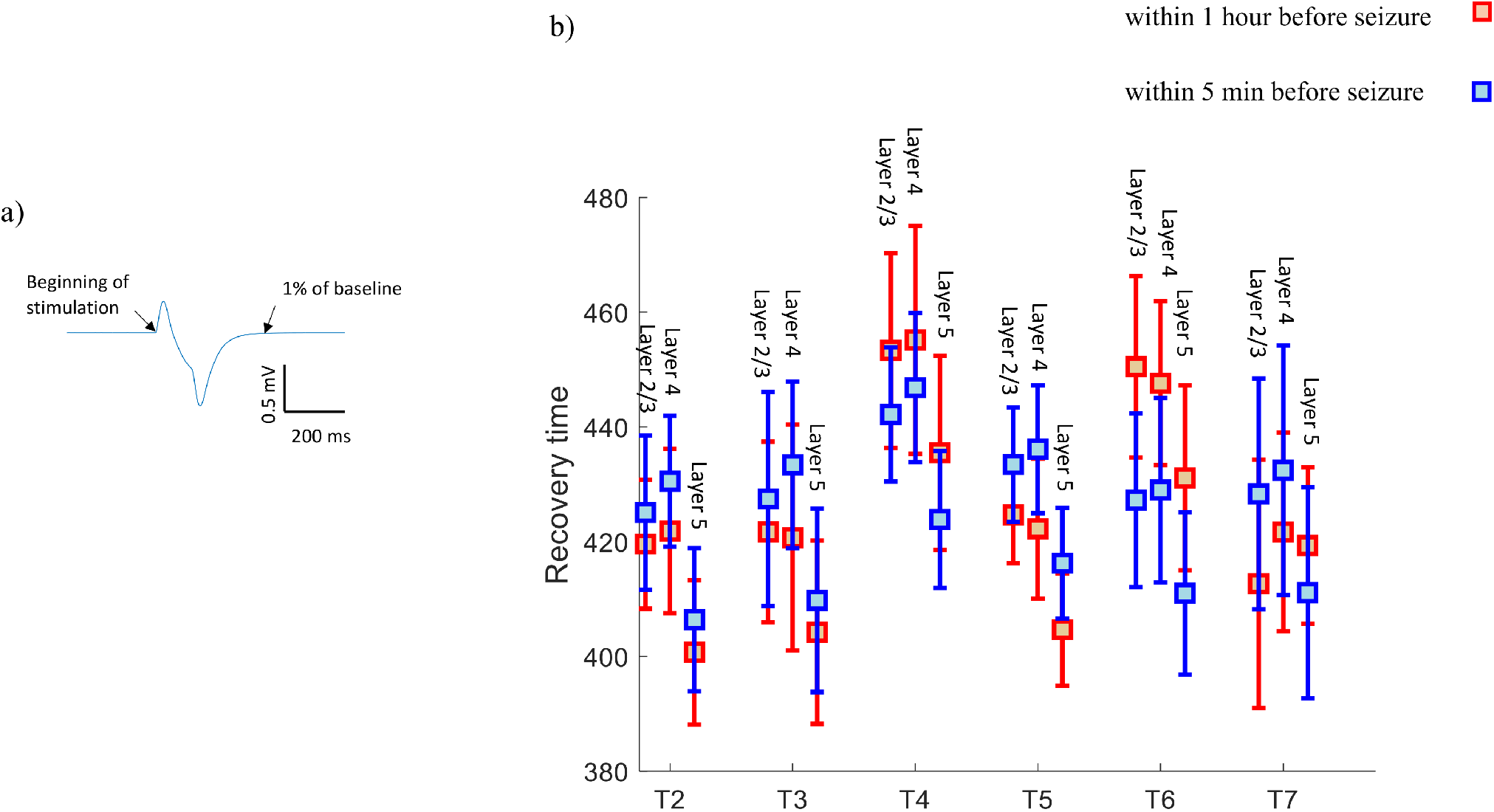
The recovery times of response to stimulation of the model fitted to the background EEG signals. a) A sample waveform of the response to of the model to stimulation. We measured the recovery time from the stimulation by measuring the time from the start of the stimulation until the response reaches to 1% of the baseline. b)The average recovery time of the model’s response to stimulation fitted to the background EEG signals within 5 mins before seizures compared to 1 hour before seizures across different rats (T2, …, T7). The error bars represent the confidence intervals. Blue represents 5 mins before seizures and red represents 1 hour before seizures.

## Discussion

In this study, we employed a neural mass model of three cortical layers to analyze background EEG recordings from a Tetanus Toxin (TT) rat model of epilepsy to investigate the relationship between neural dynamics and seizure activity. The model fitting demonstrated a close correspondence between the estimated EEG signal and the original EEG recordings, indicating estimation of the dynamics of membrane potentials and synaptic currents across various layers of the cortex.

### Correlation of the model parameters with the number of daily seizures

Correlation analyses between connectivity strengths in the model and daily seizure rates provides valuable insights into the relationship between synaptic connectivity and epileptic activity. Figures 5 and 6 illustrate the connectivity weights that exhibited statistically significant correlations with the numbers of daily seizures for different rats. The results show consistency across different rats. Table 3 summarizes the correlation results, demonstrating the prevalence of positive correlations of the strengths of Excitatory-to-Excitatory, Inhibitory-to-Excitatory, and Inhibitory-to-Inhibitory connections with the numbers of daily seizures. Notably, Excitatory-to-Inhibitory connections displayed a decrease in strength with increasing numbers of daily seizures.

We expected to observe decreases in the values of Inhibitory-to-Excitatory connections with increases in the numbers of daily seizures, which would indicate reduced inhibition in the system but, surprisingly, the positive correlation observed in Inhibitory-to-Excitatory connections contradicted this expectation. This counter-intuitive finding can be rationalized by considering the system’s response to heightened excitation through increased Excitatory-to-Excitatory and Inhibitory-to-Inhibitory connections, coupled with a reduction in Excitatory-to-Inhibitory connections. In such a scenario, an effective mechanism is required to regulate seizures and prevent an excessive surge in neural activity. The observed increase in Inhibitory-to-Excitatory connection strengths serves as a potential regulatory mechanism, contributing to the overall balance necessary to avert seizure-induced hyperexcitability and maintain neural stability.

Previous studies have shown the regulatory effect of inhibition in restraining the spread of seizures in a number of *in vivo* and *in vitro* studies [22, 26, 27, 28]. The study by Trevelyan et al. (2006) [27] demonstrated a robust inhibitory mechanism occurring before the onset of an ictal wavefront. This inhibitory process, known as feedforward inhibition, acts in opposition to or counters the propagation of epileptiform activity. In a study by Dehghani et al. (2016) [9], a plateau of high activity of inhibitory neurons during seizures was observed. This indicates that the seizures were manifested by strong ‘control’ exerted by the inhibitory neurons. A number of *in vitro* studies have shown substantial activity of inhibitory neurons before the onset of seizures, highlighting the complex interaction between excitatory and inhibitory forces in the initiation of seizures [16, 17, 31].

### Critical slowing down measure as an active biomarker

We analyzed the recorded EEG signals during the probing-on period when electrical stimulation was applied. The recovery time from electrical stimulation was measured during the periods within 5 mins before seizures compared to 1 hour before seizures. The investigation into recovery times (return to baseline after a stimuli) as a biomarker revealed interesting patterns [18, 29, 30]. Figure 8 presented recovery times in response to probing periods before seizures, showing variations across different rats. The results of this analysis indicate increases in the recovery time 5 mins before seizures compared to 1 hour before seizures in four rats (T2, T3, T6, and T7). This suggests the occurrence of critical slowing down as seizure times are approached in these rats [24].

As applying electrical stimulation directly to the brain is invasive and not always feasible, we investigate the use of the neural mass model fitted to background EEG (i.e., without probing) as a proxy measure of the critical slowing down phenomenon by applying stimulation to the computational model fitted to the background EEG 5 mins before seizures and 1 hour before seizures. We applied stimulation to both excitatory and inhibitory neurons in all layers of the cortical column and measured the recovery time from the stimulation. The results show that the recovery time after stimulation 5 mins before seizures was longer than 1 hour before seizures in four rats (T2, T3, T5, and T7). These results are consistent with the analysis of the data recorded from actual electrical stimulation to the brain, except in rats T5 and T6. Critical slowing down was observed in the analysis of the data, but this phenomenon was not observed in the results of the modeling for rat T6.

The discrepancy between the observed critical slowing down in the background EEG during the preictal period and the absence of a comparable effect in response to brain stimulation in rat T5, as well as the inverse situation for rat T6, where critical slowing down was detected in the probing data but not in the modeling results, may stem from several factors that warrant consideration. One potential factor is the inherent variability in individual neural responses, where the dynamic nature of each rat’s neural network may lead to divergent outcomes. Additionally, the limitations of the computational model employed for stimulation response, including simplifications or assumptions made in its design, could contribute to the observed disparities. Furthermore, the discrepancy might be influenced by specific physiological conditions or contextual factors that were not fully captured by the experimental or modeling approaches. Variations in a rat’s physiological state, neurochemical environment, or presence of external factors could all play roles in shaping the observed differences. These intricate interactions within the neural system highlight the complexity of studying critical slowing down and its manifestation in different experimental paradigms. To elucidate the underlying mechanisms contributing to these discrepancies, further in-depth investigations are warranted. Interdisciplinary efforts integrating experimental data and computational modeling, along with a meticulous examination of individual neural responses and potential model refinements, could offer valuable insights into the nuanced interplay between neural dynamics and the critical slowing down phenomenon. Such endeavors will contribute to a more comprehensive understanding of the complex dynamics governing epileptic seizures and enhance the translational relevance of computational models in capturing the complexities of neural behavior.

## Conclusion

In conclusion, our computational modeling approach sheds light on the intricate relationship between neural dynamics and seizure activity in the Tetanus Toxin (TT) rat model of epilepsy. By employing a neural mass model, we successfully bridged theoretical predictions with empirical observations, unraveling some mechanisms underlying epileptogenesis. Our study enhances knowledge of epilepsy, opening avenues for future research and targeted interventions in neuroscience. The insights gained not only contribute to understanding epilepsy but also emphasize the potential of computational models in advancing neuroscientific exploration.

## Supporting information

### S1. Validation of the proposed method for parameter estimation

To validate the proposed method for parameter estimation, we generated waveforms using different parameter settings applied to the neural mass model. To challenge our estimation method, we added observation noise to the waveforms with variance of 0.1. We then applied the Unscented Kalman Filter to estimate the parameters and generate the waveforms at the output of the neural mass model. Figure S1.a shows an example of an original waveform and the estimated waveform using the Unscented Kalman Filter. The estimated signal matches the original waveform very well. Figure S1.b shows the estimated parameters alongside their actual values. Despite the added noise to the waveform, the UKF was able to approximate the parameter values.

**Fig S1.**
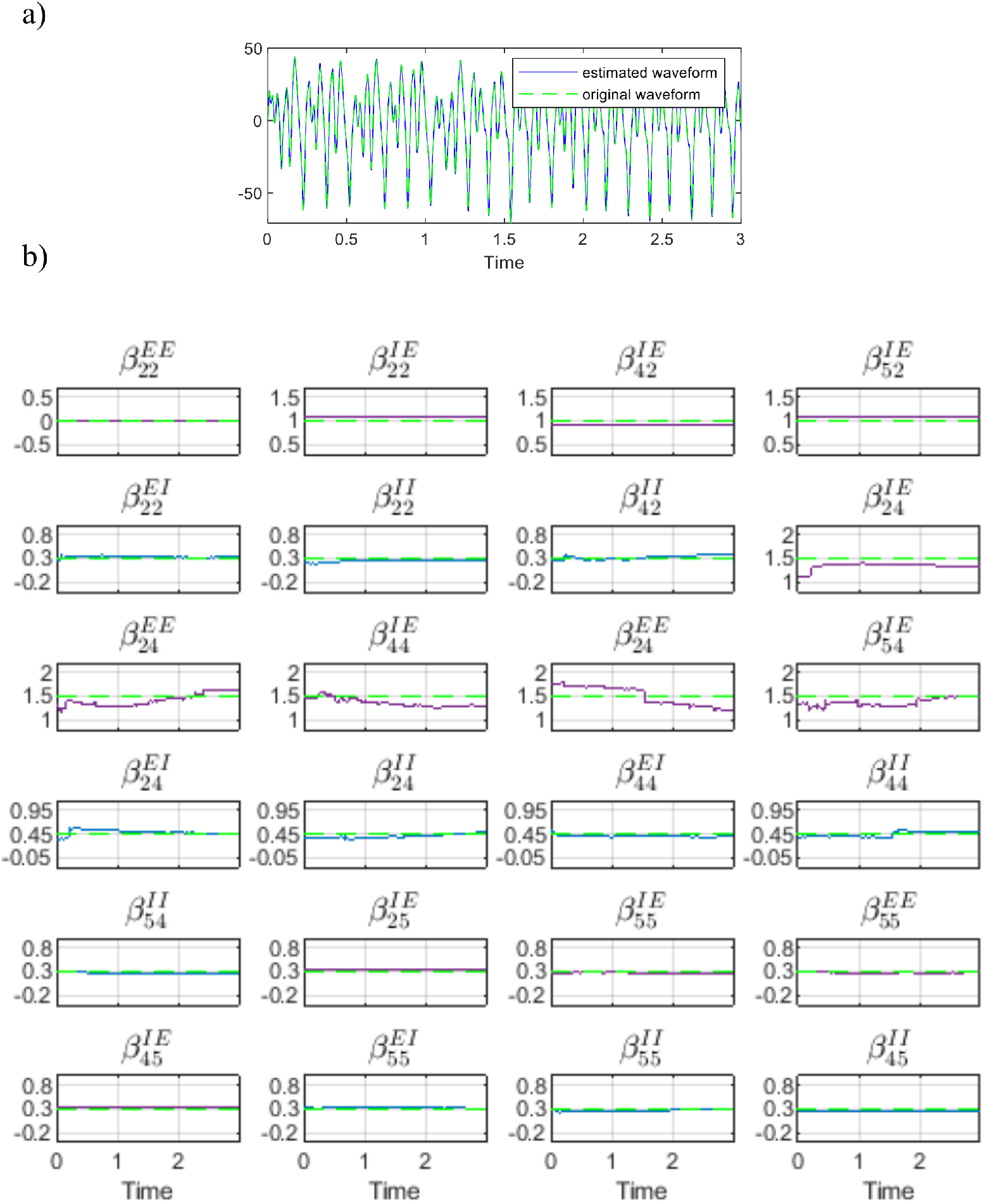
a) The estimated waveform (blue) compared to the original waveform (dashed green). b) The estimated excitatory connectivity weights (purple) and inhibitory connectivity weights (blue) compared to the original value of the parameters (dashed green).

We ran this procedure over 50 trials, and each time we set the initial values of the parameters to different random values generated by a uniform distribution between 30% added to the original parameter values. Figure S1.b displays the error in the estimated parameters (averaged over the last 2 seconds of the waveform) compared to their ground truth values across these 50 trials.

**Fig S2.**
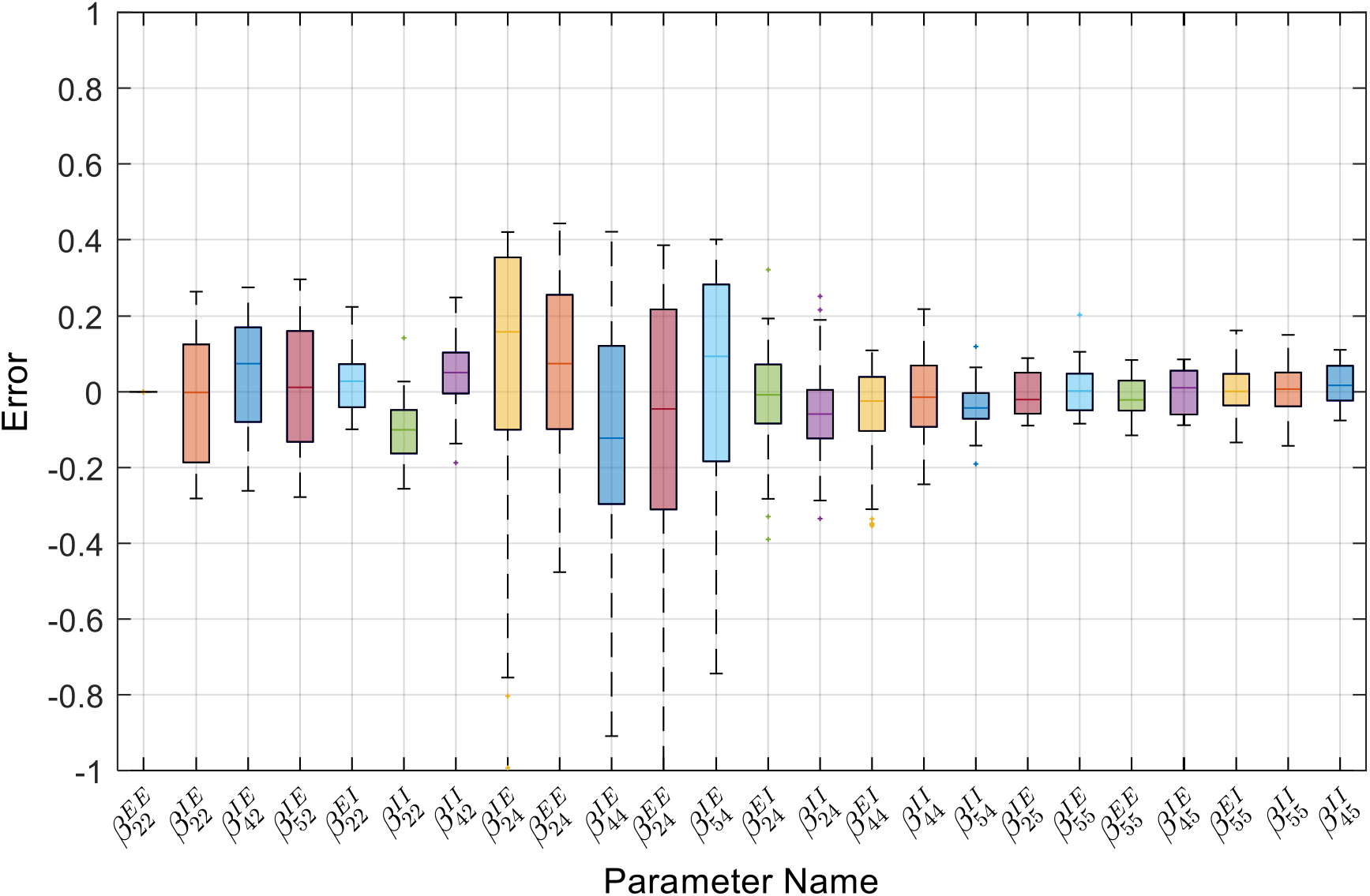
Boxplots of the errors for the estimated parameters across 50 trials of randomized parameters. The x-axis shows the parameter names while the y axis shows the error of the estimated parameters averaged over the last 2 seconds. Each boxplot represents the distribution of error for individual parameters. The box itself spans the interquartile range (IQR), with the horizontal line inside indicating the median error value. Whiskers extend to 1.5 times the interquartile range, showcasing the spread of error values within each parameter category.

## Acknowledgments

The authors thank Dr Warwick Cheung for providing the rat recordings. This work was funded by the Australian Government under the Australian Research Council’s Training Centre in Cognitive Computing for Medical Technologies (project number ICI70200030).

## Notes

### Competing Interest Statement

The authors have declared no competing interest.

### Summary of Updates

Revised. Added figures elaborating the results and modified the description of the model fitted to the signal generated by the model

